# KEDROS: A Comparative Genomics and High-Resolution Phylogenetic Profiling Workspace

**DOI:** 10.64898/2026.10.08.757521

**Authors:** Athena Marougka, Jaime Huerta-Cepas, Alexandros A. Pittis

## Abstract

Phylogenetic profiling infers gene function from the presence and absence of genes across genomes, but it loses resolution in eukaryotes, where many protein families contain several functionally distinct paralogs. Existing resources address parts of this problem separately: web services build trees for one family at a time, orthology databases provide precomputed orthologous groups for predefined species sets, and tree-profiling tools require users to supply their own trees and groupings. Here, we present KEDROS, a web platform that builds gene trees for any protein family across UniProt reference proteomes, splits each tree into subfamilies at inferred gene duplications, and profiles each subfamily separately. The organisms, the families and the taxonomic depth at which trees are split are all chosen by the user, so the same workflow serves any question that can be posed as a contrast between two sets of organisms: a pathogen and its host, a pest and non-target species, or lineages with and without a trait. KEDROS answers such contrasts at two levels. Across families and subfamilies, it ranks those present in one set and absent from the other, pointing to lineage-specific genes and candidate targets. Within a subfamily shared by both sets, it identifies amino acid positions that differ consistently between them and may underlie functional divergence, and displays them on AlphaFold structures. All analyses run in a web browser, with no installation or programming. Each of these steps exists in some form elsewhere, but to our knowledge no other resource runs them end to end, at the scale of whole proteomes, on organism sets defined by the user. We illustrate KEDROS with two cases. In the trypanothione reductase family of trypanosomatids, it recovers parasite-specific subfamilies and the substrate-binding substitutions that separate the parasite enzyme from human glutathione reductase. In the bacterial phylum Bacillota, it recovers the sporulation machinery from phenotype labels alone across 1,218 proteomes, and ranks an uncharacterized family among the strongest markers of the trait. KEDROS thus offers a general, code-free route from a comparative question to testable hypotheses about genes and residues. KEDROS is freely available at https://kedros.cgenomicslab.org.

## Introduction

Comparing gene content across genomes is an effective way to associate genes with biological functions and phenotypes. Gene trees allow such comparisons to distinguish orthologs, which diverged through speciation, from paralogs, which arose through gene duplication (Fitch 1970). Successive duplications split a gene family into subfamilies, and genes within the same subfamily are expected to share function more often than genes from different subfamilies (Gabaldón and Koonin 2013). Phylogenetic profiling builds on this by recording the presence or absence of each gene across many genomes and inferring that genes with similar profiles are functionally linked (Pellegrini et al. 1999). It has since become a general exploratory approach (Moi and Dessimoz 2023), used for questions as varied as predicting protein interactions, linking genotype to phenotype or environment, and finding drug or pesticide targets that a pathogen or pest carries but its host does not.

Despite this, phylogenetic profiling remains hard to apply to eukaryotes and hard to access for scientists without computational training. Binary profiles capture prokaryotic gene content well but lose resolution in multi-copy eukaryotic families, where a single family-level profile combines paralogs that may differ in distribution and function (Moi et al. 2020). This has motivated methods that add copy number and phylogenetic context (Dembech et al. 2023). Existing resources each cover some part of this problem. Web services such as Phylogeny.fr (Dereeper et al. 2008), NGPhylogeny.fr (Lemoine et al. 2019) and Galaxy (The Galaxy Community et al. 2024) allow tree building without coding, but for one family at a time rather than across whole genomes. Orthology and phylogeny databases such as eggNOG (Hernández-Plaza et al. 2023), OMA (Altenhoff et al. 2024), PhylomeDB (Fuentes et al. 2022) and COG (Galperin et al. 2025), together with the many others listed by the Quest for Orthologs initiative (Langschied et al. 2024), provide precomputed, genome-scale groups and trees. However, they are organized around predefined species sets and groupings and, to our knowledge, they do not offer a framework for statistically comparing groups of organisms chosen by the user. Tree-profiling tools, such as TreeProfiler (Deng et al. 2026), annotate and summarize traits along large trees, but users need to supply the trees and to define the groups to compare. Finally, running homology searches and tree inference on demand is computationally costly, and web services often become unavailable over time (Wren et al. 2017; Mangul et al. 2019).

To address these limitations, we developed KEDROS, a comparative genomics workspace that combines precomputed protein family annotations of UniProt reference proteomes with on-demand tree building in a single, code-free interface. Its central feature is high-resolution phylogenetic profiling: rather than profiling a protein family as a whole, KEDROS splits the family’s gene tree into subfamilies (clades of the tree, each corresponding to an orthologous group, OG) and profiles each subfamily separately across the chosen organisms. A discovery mode then compares two user-defined sets of organisms, for example a parasite and its hosts, and ranks the families or subfamilies present in one set and absent from the other.

In addition, because protein structure and function are encoded in sequence, positions that are conserved within groups but differ between them can point to functionally important sites (Capra and Singh 2008). When two sets of organisms share the same subfamily, KEDROS identifies the alignment positions whose amino acids differ consistently between them and displays these positions on AlphaFold protein structures (Jumper et al. 2021; Varadi et al. 2024), linking divergence in gene content and sequence to possible differences in function.

### Implementation and main features

KEDROS runs entirely on a server, so users need only a web browser, with no installation, plug-ins or programming (Fig. 1A). Analyses are organized as a workflow whose steps can also be run independently of one another: choosing organisms and protein families, building gene trees, splitting them into subfamilies, profiling these subfamilies, comparing sets of organisms, and scoring and mapping residue differences onto protein structures. Software versions, parameters and statistical details are provided in Supplementary Methods.

**Figure 1.**
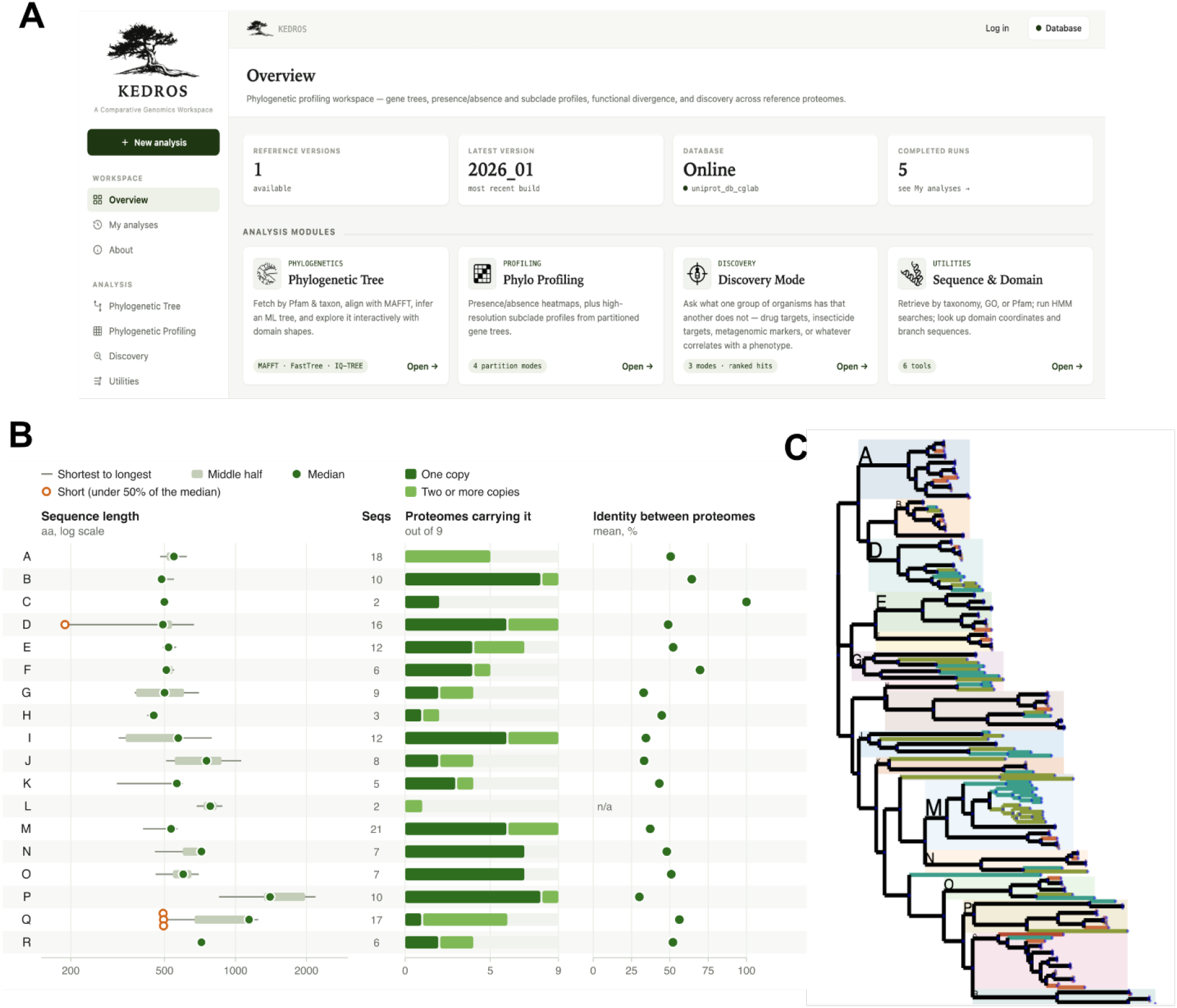
KEDROS workspace. (A) KEDROS home page and overview of functionality. (B) High-Resolution Phylogenetic Profiling tab: Subclade splitting, with Auto-duplication mode, at the level of Eukaryota (taxid: 2759) for the PF07992 protein family. The gene tree was partitioned into 18 subclades (A–R), each summarized in one row. Sequence length (amino acids, log scale): the line spans the shortest to the longest sequence in the subclade, the shaded bar marks the middle-half range, and the filled dot marks the median. Open orange circles mark sequences that are shorter than 50% of the subclade median, which may be fragments or truncated isoforms. Seqs: number of sequences in the subclade. Proteomes carrying it: number of the nine reference proteomes containing PF07992 that have at least one member of the subclade. Dark green shows proteomes with a single copy; light green shows proteomes with two or more copies (in-paralogs). Identity between proteomes: mean pairwise sequence identity (%), calculated only between copies from different proteomes. Because identity reflects both how long ago a subclade diverged and how strongly it is conserved, values should be interpreted alongside subclade depth. n/a, all members come from a single proteome. (C) The same subclades (subfamilies) displayed on the gene tree, with clades labeled.

### Precomputed protein family database

KEDROS contains the proteins of UniProt Reference Proteomes (release 2026_01) (The UniProt Consortium et al. 2025). Rather than relying on the domain annotations distributed with UniProt, all proteomes were annotated from scratch with a search against the Pfam protein family library (Mistry et al. 2021) using PyHMMER (Larralde and Zeller 2023), so that family assignments are consistent across organisms and independent of upstream annotation updates. Proteins were also assigned to KEGG orthologs with KOfam (Aramaki et al. 2020), linking them to KEGG pathways (Kanehisa et al. 2016), and Gene Ontology terms can be used to filter queries. Because all of this is precomputed, queries return quickly.

### Building gene trees on demand

For any family, users can build a gene tree from the selected proteomes in a few steps. Sequences are aligned with MAFFT (Katoh et al. 2019) or Clustal Omega (Sievers et al. 2011), gap-rich alignment columns can be removed with trimAl (Capella-Gutiérrez et al. 2009), and trees are inferred with FastTree (Price et al. 2010) or IQ-TREE (Wong et al. 2026). Trees open in an interactive explorer built on the ETE toolkit (Huerta-Cepas et al. 2016), annotated with NCBI taxonomy (Schoch et al. 2020) and protein domain architectures, and can be downloaded in standard formats, including files ready for visualization in iTOL (Letunic and Bork 2024).

### High-resolution profiling: from protein families to subfamilies

High-resolution profiling splits a family into its subfamilies before screening it. Users can define these subfamilies in four ways: by cutting the tree at a chosen distance from the root, by selecting a few sequences and taking the smallest clade that contains them, by clicking nodes in the tree explorer, or automatically, by splitting the tree at gene duplications. In the automatic mode, a node is treated as a duplication when its descendant lineages share at least one species (Huerta-Cepas et al. 2007), and the user chooses the taxonomic level at which duplications are considered (for example, Eukaryota, NCBI taxid 2759) (Fig. 1B, C). In the automatic mode, each resulting clade is an orthologous group (OG) at the chosen taxonomic level: its members descend from a single gene in the common ancestor of that taxon and are orthologs, or lineage-specific in-paralogs, of one another, whereas different clades are paralogous. Whichever mode is used, we refer to the resulting clades as subfamilies. Every subfamily is profiled separately and displayed as its own column in a heatmap aligned with the species tree, giving the presence and copy number of each subfamily rather than of the whole family. Subfamily labels are deterministic, so results can be reproduced across runs.

### Discovery mode: comparing two sets of organisms

Discovery mode compares a target set of organisms with a protected (background) set, for example a pathogen and its hosts, and ranks protein families, or the subfamilies of a split tree, by how consistently they are present in the target set and absent from the protected set. The score is the prevalence in the target set minus the prevalence in the protected set; we call it specificity, or selectivity in the target-finding modes. Each candidate is tested for differential presence with Fisher’s exact test and Benjamini–Hochberg correction for multiple testing (Benjamini and Hochberg 1995). Four preset modes cover common questions: drug targets, insecticide targets, metagenomic markers and genes correlated with a phenotype.

### Finding the residues that differ

When the two sets share a subfamily, the question turns to which amino acid positions distinguish them. KEDROS scores every alignment column for how differently the two groups use amino acids, with three complementary measures, Jensen–Shannon divergence, GroupSim and mutual information (Capra and Singh 2007; Capra and Singh 2008), and estimates significance by permutation of the group labels. Groups can be defined by clades in the tree or by taxonomy. The scores are then displayed on the AlphaFold model (Jumper et al. 2021; Varadi et al. 2024) of any protein in the alignment, in an interactive 3D viewer (Rego and Koes 2015) linked to the sequence. Before coloring, KEDROS checks that the model matches the protein’s sequence exactly, so that a different isoform or sequence version is not colored at the wrong positions, and the coloring can be exported for PyMOL.

### Use case 1: parasite-specific subfamilies and residues in a family shared with the host

This first case tests resolution inside a family: whether subfamilies confined to the parasite can be recovered from a family the host also carries, and whether the residues that differ can be located where a subfamily is shared. Trypanosomatids, the parasites that cause sleeping sickness, Chagas disease and the leishmaniases, use trypanothione and trypanothione reductase in place of the glutathione and glutathione reductase of most eukaryotes (Fairlamb et al. 1985), and trypanothione reductase has been pursued as a drug target for nearly four decades. However, the parasite enzyme is a close relative of human glutathione reductase and belongs to the same Pfam family (Pyr_redox_2, PF07992), so a family-level presence/absence screen cannot detect it. We therefore asked whether this family contains subfamilies that every trypanosomatid carries but the host lacks. We compared five trypanosomatid proteomes (*Trypanosoma brucei, T. cruzi* CL Brener, *T. cruzi marinkellei, Leishmania major* and *L. infantum*) with a protected set of *Homo sapiens, Caenorhabditis elegans* and two *Saccharomyces cerevisiae* strains.

We built the family’s gene tree (MAFFT, FastTree) and split it at duplications dated to Eukaryota, which yielded 18 subfamilies, and profiled these across the nine proteomes in drug-target mode (Fig. 2A). Three subfamilies cleared the 0.8 prevalence threshold and were absent from all four protected proteomes. Two of them, A and F, were present in all five parasites (selectivity 1.00, q = 0.071, the smallest q-value attainable when five proteomes are compared with four), while the third, R, was present in four of the five (0.80). Subfamily F is nearly single-copy, with six proteins of similar length (505–550 residues), as expected for a discrete enzyme. No protein in A or F has an informative name in UniProt, so both were found by their distribution alone. Trypanothione reductase itself falls in subfamily D, which also contains the glutathione reductases of the protected organisms, and is therefore not flagged by presence/absence; this is the situation the residue-level comparison addresses. The choice of protected set matters: with human alone, five subfamilies were returned, and adding the worm and the yeasts removed two of them.

**Figure 2.**
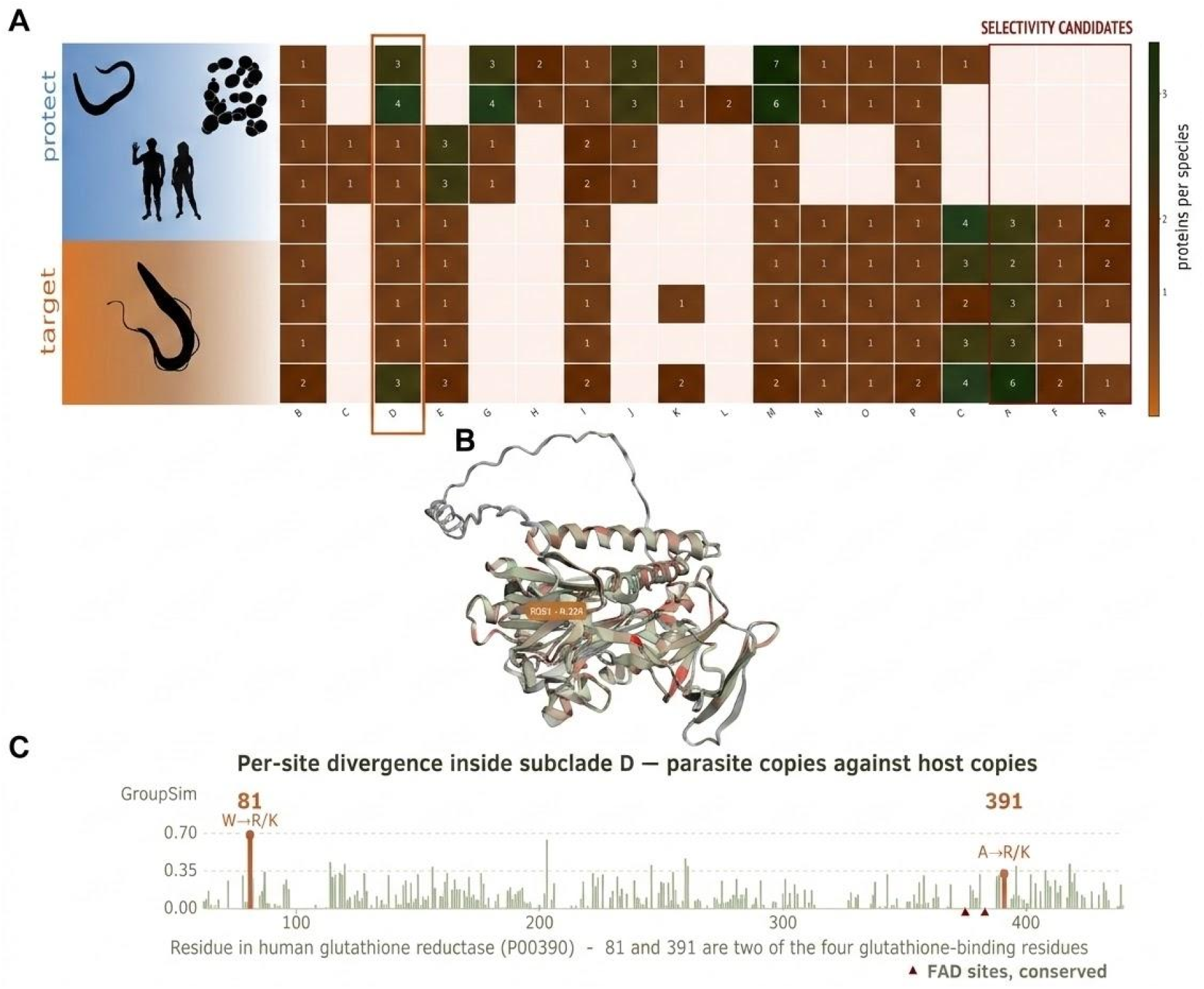
Trypanothione reductase family (PF07992) across five trypanosomatids and four protected eukaryotes. (A) Selectivity profile: proteins per species for 18 subclades. The three subclades clearing the 0.8 prevalence threshold are boxed, and subclade D, which holds trypanothione and glutathione reductase, is outlined. (B) GroupSim scores for subclade D mapped onto the AlphaFold model of human glutathione reductase (P00390). (C) GroupSim per residue of P00390: the glutathione-binding positions 81 and 391 are marked, and the FAD-binding positions (triangles) are conserved.

Subfamily D shows what KEDROS can do when a parasite and host share a subfamily: find the residues that differ. We compared the 15 sequences of subfamily D, six from the parasites (including *T. brucei* trypanothione reductase) and nine from the protected organisms (including human glutathione reductase, UniProt P00390). After removing one partial sequence of 189 residues and alignment columns with too many gaps, 348 positions remained, covering residues 62–440 of the human enzyme (Fig. 2B, C). The top-ranked position was residue 81, a tryptophan in all parasite sequences but an arginine or lysine in all host sequences, and residue 391 showed a similar contrast (alanine against arginine or lysine). Both are among the four glutathione-binding residues annotated in the human enzyme, and both are substitutions through which trypanothione reductase accommodates its larger, less acidic substrate (Bradley et al. 1991; Sullivan et al. 1991). By contrast, the other two glutathione-binding residues (74 and 158) and both FAD-binding residues (375 and 383) are conserved between the two groups. Because 161 of the 348 positions reach the same q-value (0.005), statistics alone do not single out residues 81 and 391. Their interest rests on the annotation and chemistry of these sites rather than on their rank.

### Use case 2: association with spore formation in bacteria

The second case tests the complementary situation, whole-family presence across many genomes, by asking whether a developmental program whose genetics are already known can be recovered from phenotype labels alone. Endospore formers survive an antibiotic course and germinate afterwards, which contributes to the expansion of *Clostridioides difficile* in the post-antibiotic gut (Foley et al. 2025; Müller et al. 2026), and sporulation is lost repeatedly in host-adapted gut Firmicutes in a way

In phenotype-correlation mode (minimum prevalence in the target 0.7, maximum in the background 0.3), 66 families exceeded a specificity of 0.90 (Fig. 3). In developmental order, they cover the program: initiation (Spo0A, 520/527 against 0/691), asymmetric division and engulfment (SpoIID, SpoIIE, SpoIIGA), the SpoIIIA channel that feeds the forespore (SpoIIIAB, AC, AE and AH), coat and cortex (SpoIVA, SpoIVB, YabP, YqfD), the spore core (SASP, SpoVAC/SpoVAEB, SpoVT) and germination (GerA receptor, cortex-lytic hydrolase, germination protease). The highest is SpoIID at 0.996, present in 525 of 527 spore-formers and in none of the 691 others. No family reaches 1.00, which is the expected result: sporulation is lost repeatedly within otherwise spore-forming genera (Browne et al. 2021), and a perfect score across 527 organisms would describe the taxonomy of the input rather than the biology of the trait.

**Figure 3.**
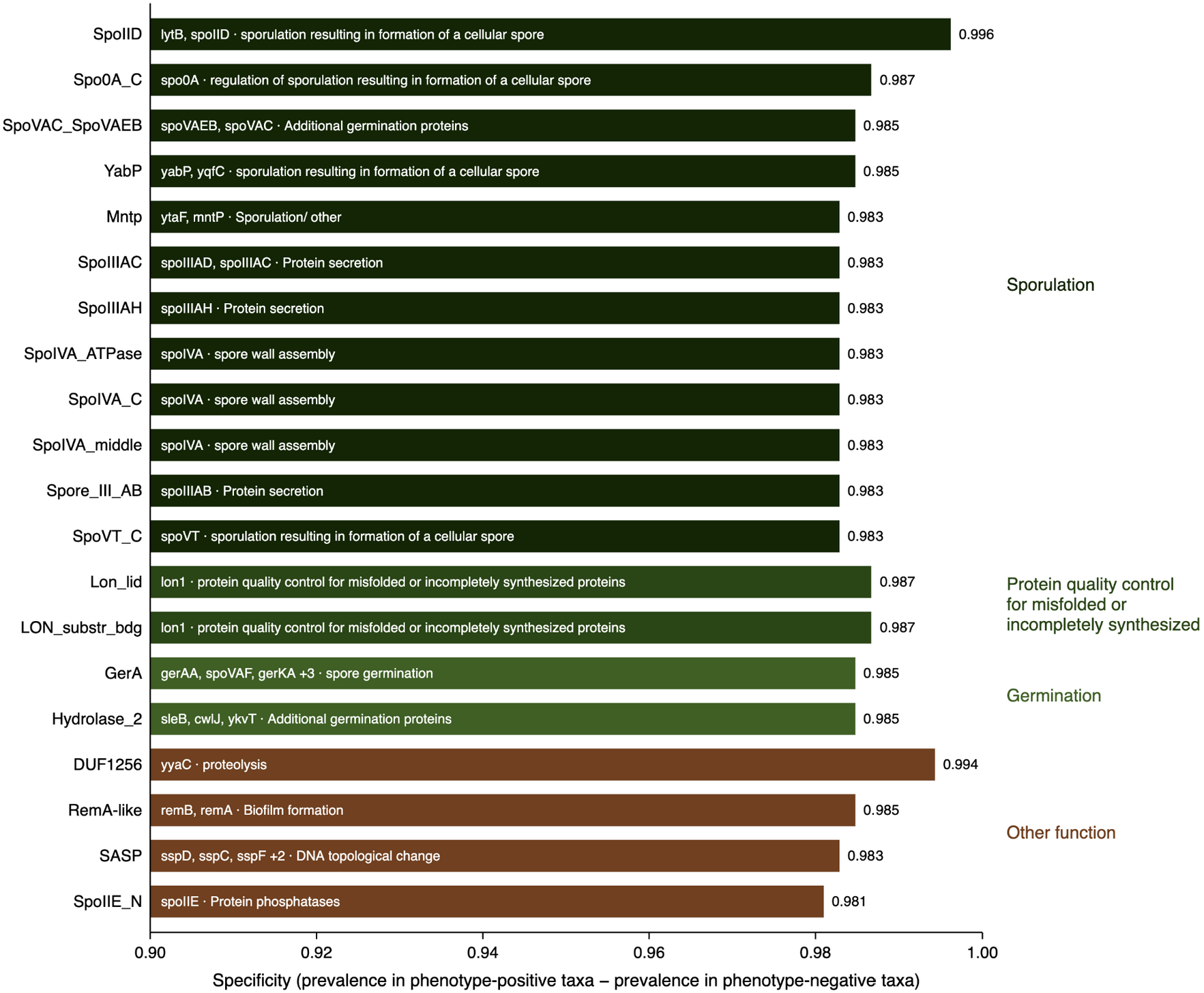
The 20 protein families most strongly associated with endospore formation, from a comparison of 527 spore-forming and 691 non-spore-forming Bacillota reference proteomes. Specificity is the prevalence of a family in the phenotype-positive set minus its prevalence in the phenotype-negative set; the value is printed at the end of each bar and the axis is truncated at 0.90. Bars are grouped by shared functional annotation (right-hand labels: sporulation, 12 families; protein quality control, 2; germination, 2; other function, 4) and ordered by specificity within each group, so the vertical order is not the overall ranking. Text inside each bar gives the *Bacillus subtilis* 168 gene names and the function, retrieved automatically from Gene Ontology (The Gene Ontology Consortium et al. 2023) biological-process terms, InterPro (Blum et al. 2025) and SubtiWiki (Pedreira et al. 2022). These 20 are the top of 300 families returned by the screen. The most specific is SpoIID at 0.996; DUF1256 (*yyaC*) is second at 0.994 but appears under “other function”, its only GO term that changes how they are transmitted (Browne et al. 2021). Asking which protein families track a trait is only useful if the method can be trusted on a trait whose molecular basis is already known, and the sporulation program of *Bacillus subtilis* is characterized stage by stage. We therefore compared seven spore-forming genera (*Bacillus, Clostridium, Paenibacillus, Geobacillus, Brevibacillus, Desulfotomaculum* and *Thermoanaerobacter*), containing 527 reference proteomes, with five non-spore-forming clades (*Staphylococcus*, Streptococcaceae, Lactobacillaceae, *Listeria* and Enterococcaceae), of 691 proteomes. Both sets lie within Bacillota and neither set is monophyletic, so the comparison is not one between two lineages, and the screen was given only the phenotype labels.

The second-ranked family is uncharacterized. DUF1256 (PF06866) is present in 524 of 527 spore-formers and in none of the 691 others (specificity 0.994), above every named sporulation gene except SpoIID. Pfam describes a family of about 180 residues restricted to Bacillales and Clostridia and distantly related to peptidase family M63, and its *B. subtilis* 168 gene is *yyaC*. A distribution screen cannot say which families are required for the phenotype and which just go with it, and that applies to DUF1256 as much as to any named gene. What it does establish is that an uncharacterized family tracks sporulation across 1,218 proteomes more tightly than most of the genes named after it.

## Conclusions

KEDROS makes phylogenetic profiling that distinguishes orthologs from paralogs available without coding: tree building, splitting trees into subfamilies, comparing sets of organisms, scoring sequence differences and mapping them onto structures run in a single interface, over a precomputed annotation of UniProt reference proteomes. The use cases show the levels at which a question can be answered. Where both sets of organisms carry a family, the answer lies among its subfamilies, and where both carry the same subfamily, among individual residues; in the trypanothione reductase family this recovered parasite-specific subfamilies that carry no informative protein name, and the substrate-binding substitutions that separate the two enzymes. Where the sets differ in whole families, presence alone is enough, and the sporulation screen recovered a developmental program from phenotype labels across 1,218 proteomes, along with an uncharacterized family that ranks above nearly all named sporulation genes. Because subfamilies are defined automatically from gene duplications, with the user setting the resolution through the taxonomic level, the approach needs no prior subfamily annotation and extends to any two sets of organisms a user can define. It cannot separate families required for a phenotype from those that merely share its history. It returns specific, testable hypotheses: candidate drug and pesticide targets, clade markers, and genes associated with a trait.

## Acknowledgments

We thank the members of the Comparative Genomics Lab (CGLab) at IMBB-FORTH for testing the public beta and for feedback on the interface. KEDROS is hosted at IMBB-FORTH.

## Supplementary material

Supplementary material is available online with this version of the preprint.

## Data availability

KEDROS is freely available without registration at https://kedros.cgenomicslab.org/ (public beta, v0.9.0) and runs in all major web browsers. It is implemented in Python with the Django web framework and a MySQL database.

