## Supplementary Material for "KEDROS: A Comparative Genomics and High-Resolution Phylogenetic Profiling Workspace"

References cited in both the main text and this file are listed in the main manuscript; those cited only here are listed under Supplementary References at the end of this file.

In this supplementary file, we present in more detail the methods used by KEDROS, followed by two additional use cases (Use cases 3 and 4). The methods apply to every analysis in the main text and supplementary use cases. Software versions are the ones installed on the public server (Table M1).

### Supplementary Methods

#### 1. Implementation and software

KEDROS is a web application developed in Python 3.11, with the Django framework. It is served by Gunicorn behind an nginx web server and reads the data from a MySQL 8.0 database holding 34,230 UniProt reference proteomes, and their precomputed annotations. The analyses run on the server: the browser sends the request, the server runs the external tools listed in Table M1, and the results are returned as interactive objects or downloadable files in standard formats. Long jobs run in the background. Interactive trees are visualized with ETE Toolkit version 4 (ETE4) tree explorer, and NCBI Taxonomy is accessed through the ETE4 NCBITaxa module.

**Table M1.** Software and data resources used by KEDROS.

| Purpose | Software or resource | Version | Reference |
|---|---|---|---|
| Protein sequences and proteomes | UniProt Reference Proteomes | release 2026_01 | UniProt Consortium 2025 |
| Protein family annotation | Pfam-A HMM library (INTERPRO) | 27,481 families | Mistry et al. 2021 |
| HMM search | PyHMMER (HMMER3) | 0.12.3 | Larralde and Zeller 2023 |
| KEGG orthologs | KOfam HMM profiles | 27,816 profiles | Aramaki et al. 2020 |
| Gene Ontology terms | UniProt GO annotations | release 2026_01 | Gene Ontology Consortium 2023 |
| Taxonomy | NCBI Taxonomy, through ETE4 NCBITaxa | — | Schoch et al. 2020 |
| Multiple sequence alignment | MAFFT (FFT-NS-2 and E-INS-i) | 7.505 | (Katoh et al. 2019) |
| Multiple sequence alignment | Clustal Omega | 1.2.4 | Sievers et al. 2011 |
| Alignment trimming | trimAl | 1.5.1 | Capella-Gutiérrez et al. 2009 |
| Tree inference (fast) | FastTree | 2.2.0 | Price et al. 2010 |
| Tree inference (maximum likelihood) | IQ-TREE | 3.1.1 | Wong et al. 2026 |
| Tree handling and interactive explorer | ETE Toolkit (ETE4) | 4.3.0 | Huerta-Cepas et al. 2016 |
| Protein structures | AlphaFold Protein Structure Database (API) | current release | Jumper et al. 2021; Varadi et al. 2024 |

| | | | |
|---|---|---|---|
| 3D structure viewer | 3Dmol.js | — | Rego and Koes 2015 |
| Functional labels for Discovery hits | InterPro, UniProt and SubtiWiki web services | — | Blum et al. 2025; Pedreira et al. 2022 |
| Web application | Django; MySQL | 5.2; 8.0 | — |

### 2. Precomputed protein family database annotations

The database contains the 2026_01 release UniProt Reference Proteomes, consisting of 133,566,197 protein accessions, and 33,508 organisms, with accession ids, names, organisms, sequences and Gene Ontology terms and Protein Family (Pfam) annotations as distributed by UniProt. Pfam annotations were also computed by us (27,481 hmm profiles), with PyHMMER (HMMER 3), with the hmmsearch function, using the Pfam gathering thresholds (bit_cutoffs = "gathering"), so the assignment of proteins to families is consistent across all organisms. For every hit the database stores the full-sequence and per-domain E-values and scores and the domain coordinates, which are used for domain architectures and, optionally, to apply a stricter E-value cutoff when sequences are retrieved from the database.

Proteins of 55 reference proteomes were also assigned to KEGG orthologs by searching the KOfam profile library (27,816 profiles) with the same PyHMMER pipeline, and linked to KEGG pathways (48,863 ortholog-to-pathway links across 503 pathways).

Organisms are selected through NCBI taxonomy. A taxon ID given by the user is expanded to all of its descendants, including intermediate nodes. This matters because reference proteomes are attached to species taxon IDs, and in NCBI taxonomy a species is often an internal node with sequenced strains below it. Expanding to leaves only would miss exactly those species. The expanded list is then intersected with the organisms present in the database, and the page reports both numbers.

### 3. Building gene trees

For a gene tree, KEDROS retrieves every protein of the selected proteomes that carries the chosen Pfam family. Several families can be combined, requiring either any of them or all of them in the same protein. Users can exclude taxa, apply an E-value cutoff, limit the number of proteins per proteome, or add their own sequences from an uploaded FASTA file, which are merged with the database sequences. The sequence name format is {taxonID.accession}, so that each leaf of the tree can be assigned to its organism downstream. The sequences are then aligned, trimmed and used to infer the tree. The tools used for the tree building are described in Table M2.

**Table M2**. Alignment, trimming and tree-inference steps, with the options available in KEDROS. Defaults are shown in bold. N is the number of CPU threads.

| Step | Option | Command run by KEDROS | Notes |
|---|---|---|---|
| Alignment | MAFFT | mafft --anysymbol --thread N | Default strategy (FFT-NS-2). --anysymbol accepts selenocysteine (U) and pyrrolysine (O), which occur in UniProt sequences. |
| | MAFFT E-INS-i | einsi --anysymbol --thread N | Slower, iterative; for sequences with long unaligned regions. |
| | Clustal Omega | clustalo --threads N -i in -o out | Default settings. |
| Trimming | trimAl | trimal -gt 0.01 | Removes alignment columns in which fewer than 1% of sequences have a residue. The threshold can be changed by the user. |
| Tree inference | FastTree | fasttree -lg | Approximately maximum-likelihood tree under the LG model (Le and Gascuel 2008) with CAT rate categories; branch supports are SH-like local supports. |
| | IQ-TREE | iqtree -s aln -mset LG -B 1000 -T N | Model selected with ModelFinder (Kalyaanamoorthy et al. 2017) among LG-based models; 1,000 ultrafast bootstrap replicates (Hoang et al. 2018). |

Trees are visualized in the ETE4 tree explorer. For display, the tree is rooted at its midpoint and multifurcations are resolved. Leaves are annotated with NCBI taxonomy and with the Pfam domain architecture of each protein, drawn from the stored domain coordinates. Trees, alignments and sequences can be downloaded in standard formats (Newick, FASTA), together with annotation files for iTOL (Letunic and Bork 2024). For high-resolution profiling, the tree is displayed with the root written by the inference program; the root used for the automatic split can be chosen separately (Section 4). A multifurcation at the root is resolved and the tree is ladderized, and this exact topology is saved, so that the explorer and the subclade partition always refer to the same tree

### 4. High-resolution profiling: splitting a family into subfamilies

High-resolution profiling splits a gene tree into subclades, each corresponding to one subfamily, and profiles every subclade separately. The four ways to define subclades are summarized in Table M3. For any mode chosen, subclades are labeled A, B, C, … in the order in which they appear in the ladderized tree, from top to bottom, so that the same tree always gives the same labels and the columns of the profile are reproducible.

**Table M3**. The four ways to define subclades.

| Mode | How subclades are defined | Parameters |
|---|---|---|
| Distance from the root | The tree is cut at a chosen distance from the root (sum of branch lengths). Every branch crossing that distance starts a subclade. | Cut distance (slider, from 0 to the longest root-to-leaf distance). |
| Common ancestor of chosen sequences | The user lists groups of sequences; each subclade is the smallest clade containing a group. | Sequence lists. Leaves outside every chosen clade are grouped as "unassigned". |
| Nodes picked in the explorer | The user clicks nodes in the tree explorer; each picked node defines a subclade. | Picked nodes. Leaves outside every picked node are left out. |

| | | |
|---|---|---|
| Automatic, at gene duplications | The tree is split at gene duplications dated to a chosen taxonomic level (see text). | Taxonomic level (NCBI taxon ID, e.g. Eukaryota, 2759). Minimum of 2 species on each side of a duplication; subclades with fewer than 2 sequences are grouped as "unassigned".<br>Root: as written by the tree program (default) or midpoint. |

**Automatic splitting at duplications**. Only the species that belong to the chosen taxonomic level are considered. Following the species-overlap approach (Huerta-Cepas et al. 2007), an internal node of the gene tree is treated as a duplication when the same species occurs below two of its child branches. A recent expansion inside a single species also satisfies this rule, and splitting at such nodes would break the tree into many very small groups. For this purpose, KEDROS keeps a duplication only if at least two of its child branches each contain at least two different species. The duplication is then dated: the most recent common ancestor of the species below the node is found in NCBI taxonomy, and the duplication is used as a subclade boundary only if this ancestor is the chosen taxonomic level or older. Duplications that happened inside the chosen level are lineage-specific expansions and do not split the tree. Starting from the root, the tree is split at every such duplication and the descent continues into each side for as long as another dated duplication lies below it (up to eight nested levels). The taxonomic level therefore sets the resolution: choosing an older level gives fewer, broader subfamilies, and choosing a younger level gives more, narrower ones.

**Rooting.** Because duplications are read from the root of the gene tree downwards, the position of the root affects the split. FastTree and IQ-TREE place the root arbitrarily, so before splitting, the user chooses how the tree is rooted: as written by the tree-inference program (the default) or at its midpoint, and KEDROS reports the root used and the subclades on each side of it. The split is computed on a re-rooted copy of the tree, so the tree shown in the explorer is not changed. For the PF07992 tree of Use case 1 (Fig. 1B), the root written by FastTree is one of the roots with the fewest inferred duplications (67, the minimum over all possible roots). Re-rooting the same tree at its midpoint gives 17 subclades instead of 18: sixteen are identical, subclades B and C merge into one, and subclades A, D and F, on which Use case 1 relies, are unchanged.

**Profile**. For each subclade and proteome, KEDROS counts the proteins of the subclade that the proteome contains. The result is shown as a heatmap with one column per subclade and one row per proteome, ordered by the NCBI taxonomy tree of the selected organisms, so that the presence and copy number of every subfamily can be read across the species. The same matrix is the input of Discovery mode at the subclade level (Section 6).

### 5. Subclade statistics

For any partition, KEDROS can summarize each subclade (Fig. 1B of the main text). These statistics are descriptive and are meant to characterize a subclade, for example one selected by a Discovery screen. They are not tested and not ranked, because the members of a subclade share ancestry and are not independent observations, and because the number of proteomes is usually small. The measures are listed in Table M4.

**Table M4.** Measures reported for each subclade.

| Measure | Definition |
|---|---|
| Sequences | Number of sequences in the subclade. |
| Proteomes carrying it | Number of proteomes in the tree with at least one member, split into proteomes with one copy and with two or more copies (in-paralogs). The total is the number of proteomes in the tree, i.e. those carrying the family. |
| Sequence length | Shortest, longest, median and middle-half range (interquartile range) of the full, unaligned sequence lengths, in amino acids. |
| Short sequences | Sequences shorter than 50% of the subclade median length, flagged only when the subclade has at least 3 sequences. These may be fragments or genuinely shorter forms; the rule cannot tell them apart. |
| Depth | Mean branch-length distance from the common ancestor of the subclade to each of its members. |
| Identity between proteomes | Mean pairwise sequence identity, computed only between copies from different proteomes (details in text). Not available when all members come from one proteome. |

**Identity between proteomes.** Identity is calculated from the untrimmed alignment. For each pair of sequences, it is the number of identical residues divided by the number of columns where both sequences have a residue. Gaps and unknown residues are ignored. A pair is used only if the two sequences share at least 30 columns and at least 50% of the residues of the shorter sequence, so that a fragment overlapping a small part of a full-length protein cannot appear almost identical to it. The average of identities is first calculated within each pair of proteomes and then across proteome pairs, each pair of proteomes weighted equally, so that recent copies within one proteome (in-paralogs) cannot dominate the value. When a subclade has more than 20,000 sequence pairs, a random sample of pairs is used (with a fixed seed), and this is reported. Because identity falls both with the age of a subclade and with weaker constraint, it should be read together with depth.

### 6. Discovery mode: comparing two sets of organisms

Discovery mode compares a target set of organisms with a background set (the protected set when the background must not be harmed, as for drug and pesticide targets). It can be run on whole protein families, using the precomputed annotation of all proteomes in the two sets, or on the subclades of a high-resolution profile (Section 4).

**Scores.** For each family or subclade, KEDROS counts the organisms of each set that carry it, and calculates its prevalence in the target set (pT, the fraction of target organisms carrying it) and in the background (pB). The specificity, or selectivity, is $pT - pB$, from 1 (in every target organism and in no background organism) to $-1$. A candidate is called when pT is at least the minimum prevalence in the target and pB is at most the maximum prevalence in the background. Each mode has its own default thresholds and its own ranking (Table M5). Raw counts are always shown together with the fractions. Copy number can also be used for ranking: the copy-number enrichment is (mean copies per target organism + 1) / (mean copies per background organism + 1).

**Statistical test**. For each candidate, the $2 \times 2$ table of target and background organisms with and without it is tested with a one-sided Fisher's exact test for enrichment in the target set, and p-values are corrected for multiple testing with the Benjamini–Hochberg procedure (Benjamini and Hochberg 1995). For whole families, the correction is over the families that pass the two thresholds. For subclades, every subclade is scored and the correction is over all of them. Before a screen is run, KEDROS also reports the smallest p-value the comparison can reach, which is the p-value of a perfect separation and depends only on the sizes of the two sets. For example, one target organism against six protected organisms can never give p below 0.14, however clean the result, and the user is told this before running the screen.

**Table M5.** The four Discovery modes. All modes use the same scores and test; they differ in default thresholds and in how candidates are ranked.

| Mode | Question | Default thresholds (min. in target / max. in background) | Ranking |
|---|---|---|---|
| Drug target | Present in the pathogen, absent from the host | 0.8 / 0.0 | Selectivity ($p$T − $p$B) |
| Insecticide target | Present in the pest, absent from species to protect | 0.8 / 0.0 | Selectivity ($p$T − $p$B) |
| Metagenomic marker | Present in nearly every member of a clade, rare elsewhere | 0.9 / 0.05 | Coverage of the clade ($p$T) first, then cleanliness (1 − $p$B) |
| Phenotype correlation | Presence tracks a trait, in either direction | 0.7 / 0.3 | Size of the difference $\lvert p\mathrm{T} - p\mathrm{B}\rvert$; when every subclade is ranked, one depleted in the target ranks as high as one enriched |

**Interpreting the test.** There are two limits in the test that should be considered. First, reference proteomes are a census of the chosen organisms, not a random sample. Second, related organisms are not independent: a family inherited once by the common ancestor of five mites is counted five times, which makes Fisher's test overconfident. For drug, insecticide and marker screens the decision is therefore made by the thresholds – a single protected organism carrying a candidate rules it out, whatever its p-value – and the q-value is reported as supporting information. Phenotype correlation mode asks whether an association could arise by chance, and it is the one where the test carries most weight.

**Organisms in the comparison.** At the subclade level, only organisms that are rows of the profile are counted, so prevalences are fractions of what was measured. An organism listed in both sets is counted only in the protected set. Profiled organisms in neither set are shown as outgroups: they do not take part in the statistics, but they show whether a subclade absent from the protected set is truly restricted to the target or simply was not sampled elsewhere.

**Functional labels.** In the metagenomic-marker and phenotype modes, the top hits are given a short functional label for the bar chart (as in Fig. 3 of the main text). Labels come, in order of preference, from Gene Ontology biological-process terms of the target proteins carrying the family (from the local database), from the InterPro entry of the Pfam family and its GO terms, from the gene names of reviewed UniProt entries in the target set, and, for Bacillus subtilis genes without GO terms, from SubtiWiki categories. Hits are grouped by the GO terms shared by the most families.

### 7. Finding the residues that differ between groups

When two sets of sequences share a subfamily, KEDROS scores every column of their alignment for how differently the two groups use amino acids. The two groups can be defined by clades in the tree or by taxonomy. Three complementary scores are calculated (Table M6). Sequences are weighted with position-based weights (Henikoff and Henikoff 1994), so that a group of many closely related sequences does not dominate a column.

**Table M6.** Per-column scores used to find residues that differ between two groups.

| Score | What it measures | Calculation | Gaps |
|---|---|---|---|
| Jensen–Shannon divergence (JSD) | Conservation of a column across the whole alignment; needs no groups | Divergence between the column's amino-acid distribution and the BLOSUM62 background distribution (Henikoff and Henikoff 1992), smoothed with the neighboring ±3 columns (weight 0.5) (Capra and Singh 2007) | Excluded, then the score is multiplied by the fraction of sequences with a residue |
| GroupSim | Columns conserved within each group but different between groups (specificity-determining positions) | Mean residue similarity within groups minus mean similarity between groups, with BLOSUM62 rescaled to 0–1 (Capra and Singh 2008) | Column not scored if any group has more than 10% gaps |
| Mutual information (MI) | How well the residue in a column predicts the group | Mutual information between residue and group label, corrected for small-sample bias (Buslje et al. 2009) and divided by the entropy of the group labels, so 1 means the residue identifies the group with certainty | Treated as a 21st symbol, so an insertion in one group counts as a difference |

**Significance.** Significance is estimated by permutation. The group labels are shuffled among the sequences (200 times by default ; up to 2,000 can be chosen), all columns are scored again each time, and the p-value of a column is (number of permutations reaching the observed score + 1) / (number of permutations + 1). The smallest possible p-value is therefore 1 / (number of permutations + 1), which is 0.005 with 200 permutations. p-values are corrected across columns with the Benjamini–Hochberg procedure. When many columns reach this smallest possible value, as in Use case 1, the test cannot rank them further and more permutations are needed to separate them. KEDROS raises a warning when a group has fewer than four sequences or when the groups cover less than 25% of the alignment, because group scores computed from very few sequences can be unreliable.

**Mapping the scores onto protein structures.** The column scores can be displayed on the structure of any protein in the alignment. The predicted structure is retrieved from the AlphaFold Protein Structure Database through its API, using the protein's UniProt accession. A model is used only if its accession matches and its sequence is identical to the sequence that was scored, otherwise nothing is colored and the user is informed with a message on the page. This prevents scores from being placed on the wrong residues of a different isoform or sequence version. Residue numbers are taken from the untrimmed alignment, counting every residue of the chosen sequence, because trimming removes columns and would shift the numbering. Structures are displayed with 3Dmol.js, linked to the sequence, and the coloring can be exported as a PyMOL script.

### Supplementary Use Cases

Two additional use cases follow: Use case 3, a metagenomic marker for Bifidobacterium in human stool samples, and Use case 4, a target for an acaricide that spares bees. They use the methods described above with the parameters given in each case.

### Use case 3: Metagenomic marker - detecting Bifidobacterium in human samples

This supplementary case detects metagenomic markers from bacteria that could be found in human stool samples. Antibiotic-driven loss of beneficial gut bacteria is a central concern in treatment (Abdel-Hady et al. 2026; Müller et al. 2026). Bifidobacterium is the genus most consistently reported to be affected by antibiotics. Measuring it depends on detecting the genus specifically in a mixed sample, and 16S rRNA is a poor answer at genus resolution because copy number varies, and close relatives are near-identical. We therefore asked which protein family, if any, is carried by essentially every Bifidobacterium and by almost nothing else in the background. A marker is a whole family that one clade carries and others do not, so no partitioning is needed.

The clade to detect was *Bifidobacterium* (NCBI taxid 1678), which expands to 921 taxa, of which 97 have a reference proteome in UniProt knowledgebase release 2026_01. The screen ran in metagenomic marker discovery mode of KEDROS, with a minimum share of the clade of 0.9 and a maximum share of the background of 0.05, against two backgrounds. The first was a gut community: thirteen families (Enterobacteriaceae, Bacteroidaceae, Lachnospiraceae, Oscillospiraceae, Clostridiaceae, Erysipelotrichaceae, Veillonellaceae, Streptococcaceae, Lactobacillaceae, Enterococcaceae, Akkermansiaceae, Coriobacteriaceae and Fusobacteriaceae), resolving to 1,484 proteomes. The second included everything else, consisting of five bacterial phyla plus the whole phylum of Actinomycetota decomposed into the 58 clades that do not contain Bifidobacterium, resolving to 15,259 proteomes with no organism present in both groups at the same time. That deliberately keeps Gardnerella, Scardovia, Alloscardovia, Parascardovia and the other Bifidobacteriaceae in the background, so a marker has to separate the genus from its own family.

Against the gut community, 66 families pass both thresholds, and nine reach a perfect score. Against the full background, of the 3,516 protein families present in the genus at all, 11 pass and none of them is perfect (Fig. S1, Table S1). The best hit is DUF6466 (PF20070), in 96 of 97 Bifidobacterium proteomes at one copy each, and in 22 of 15,259 other bacteria, including the genera of its own family. The three markers that are commonly used were all declined, each for a different reason. Fructose-6-phosphate phosphoketolase defines the bifid shunt and is the classical biochemical test, but its family (PF03894) occurs in 10,191 taxa. The activity is documented in intestinal Lactobacillus (Bolado-Martínez et al. 2012), which is in the gut background, and in Alloscardovia within Bifidobacteriaceae itself (Sechovcová et al. 2017). It is diagnostic of the genus's metabolism, not of its gene content. On the other hand, 16S rRNA is not a protein and cannot be expected to appear in a protein-domain database, and groEL is a universal chaperonin carried by every organism in the background. Both work because their sequence varies between clades, not their presence.

What the screen cannot predict is whether DUF6466 is expressed, surface-exposed in practice, or conserved enough at the nucleotide level for a primer pair: distribution is not function, and presence is not abundance. What it establishes is that from gene content alone, with no annotation supplied, it recovered a family that Pfam had independently flagged as bifidobacterial and probably surface-located, while declining the three markers already in use. Whether DUF6466 outperforms 16S in a stool sample is a question for the wet lab.

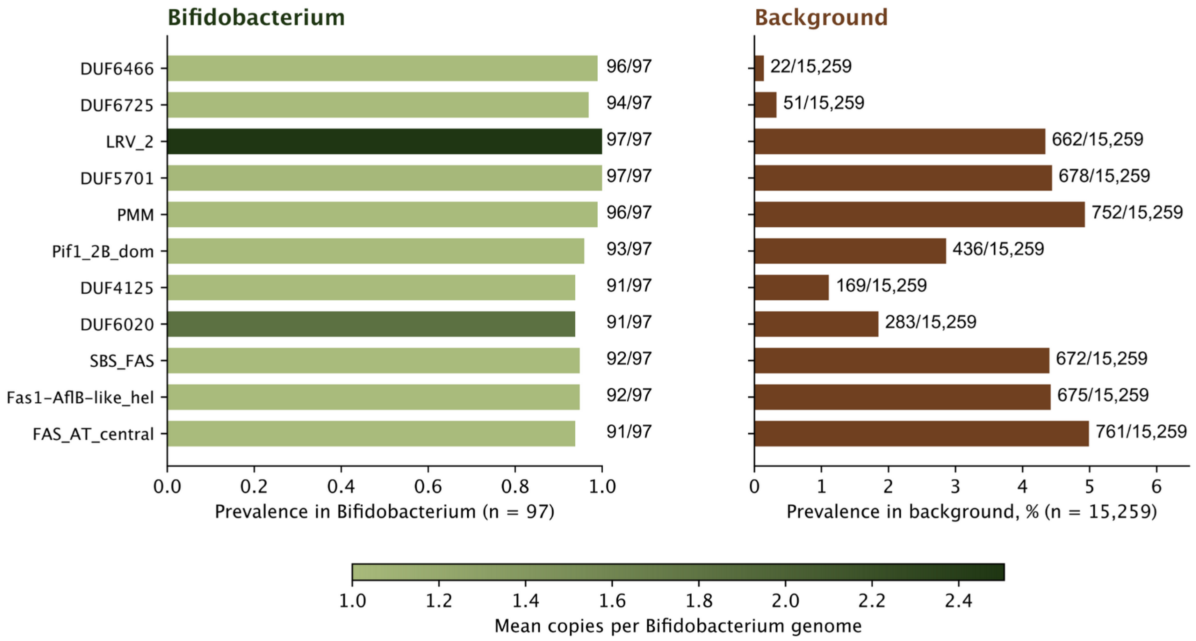


**Figure S1.** The 11 protein families carried by nearly every Bifidobacterium reference proteome, and by almost no other bacterium. **Left**: prevalence across the 97 Bifidobacterium proteomes, annotated by the mean number of copies per genome. **Right**: prevalence across the 15,259 proteomes of the background, as a percentage. Counts beside each bar are the proteomes carrying the family. Families are ordered by marker score.

**Table S1.** The five most specific of the 11 families passing the full background. Marker score is the prevalence in the clade minus the prevalence in the background: q is the Benjamini-Hochberg corrected Fisher p-value. The full table can be found in supplementary_case_3.tsv.

| Family | In Bifidobacterium | In background | Marker score | q |
|---|---|---|---|---|
| **DUF6466 (PF20070)** | 96 / 97 | 22 / 15,259 | 0.988 | $7.8 \times 10^{-226}$ |
| DUF6725 (PF20486) | 94 / 97 | 51 / 15,259 | 0.966 | $1.7 \times 10^{-202}$ |
| LRV_2 (PF25591) | 97 / 97 | 662 / 15,259 | 0.957 | $8.4 \times 10^{-130}$ |
| DUF5701 (PF18959) | 97 / 97 | 678 / 15,259 | 0.956 | $6.3 \times 10^{-129}$ |
| PMM (PF03332) | 96 / 97 | 752 / 15,259 | 0.940 | $1.1 \times 10^{-121}$ |

### Use case 4: Insecticide targeting - An acaricide that spares bees

In this supplementary case, we used KEDROS to find targets for an acaricide that would not harm bees and other animals , but would only target acari. Amitraz is the most common acaricide against Varroa destructor, the parasitic mite of *Apis mellifera*, and it is effective, because the mite's β-adrenergic-like octopamine receptor is more sensitive to it than the bee's (Guo et al. 2021). However a Y215H substitution in the same receptor is associated with contemporary amitraz resistance in the field

(Rinkevich et al. 2023). A replacement target has to be present in the mite and absent from the bee. We therefore asked whether ionotropic glutamate receptors (iGluRs) contain a group like this, since the arthropod neuromuscular junction (NMJ) is glutamatergic and the vertebrate one cholinergic. Every animal carries iGluRs, so a family-level screen does not give an answer. The question only exists at the level of subfamilies.

We performed high-resolution phylogenetic profiling for both the ANF_receptor (PF01094) and Lig_chan (PF00060) domains, for 17 reference proteomes, giving 219 sequences, aligned with MAFFT, trimmed with trimAl and built into a tree with FastTree. The tree was split at duplications dated to Arthropoda (tax id 6656), yielding 14 subfamilies, which were profiled in insecticide-target discovery mode of KEDROS (minimum prevalence in the target 0.8, maximum in the protected set 0.0). The target set was five mites and ticks (Varroa destructor, Tetranychus urticae, Sarcoptes scabiei, Ixodes scapularis and Rhipicephalus microplus), while the protected set was six species that must not be harmed, two bees (Apis mellifera and Bombus terrestris) and four vertebrates (Homo sapiens, Mus musculus, Bos taurus and Danio rerio). Six animals - three insects, Caenorhabditis elegans, Lottia gigantea and Capitella teleta - were profiled as outgroups, meaning they are in the set but not tested, because their distribution is what distinguishes a group genuinely restricted to mites from one that is unsampled elsewhere.

Two subfamilies are carried by mites and ticks and by no protected species (Fig. S2, Table S2). Subclade D is present in all five arachnids, Varroa included, at one to three copies each, and in none of the six protected species (selectivity 1.00, $q = 0.030$). Subclade A is in four out of five, at one to six copies, and likewise in none of the protected species (0.80, $q = 0.106$). Both are absent from every vertebrate, so a compound directed at either starts from the separation that amitraz is losing. Neither can be an answer on its own: with five target species, a group present in all of them reaches $q = 0.030$ at best, so these are candidates ranked by distribution, not established targets.

Subclade C holds the characterized Drosophila neuromuscular-junction subunits, which is where an arthropod-selective glutamatergic target would be looked for first, but both bees carry it and no arachnid does. Choosing it would spare the mite and hit the pollinator. The screen was given no annotation, only the distribution of subfamilies across the seventeen proteomes, and it placed the receptors the literature mentions on the wrong side of the comparison from the outset.

What this cannot establish is that subclade D is druggable, expressed at the mite neuromuscular junction, or distinct enough in its ligand-binding pocket for a compound to discriminate. What it does establish is that inside a receptor superfamily every animal shares, gene content alone separates two groups the mites have and the bees do not, which is the selectivity window an acaricide needs and the one amitraz is losing.

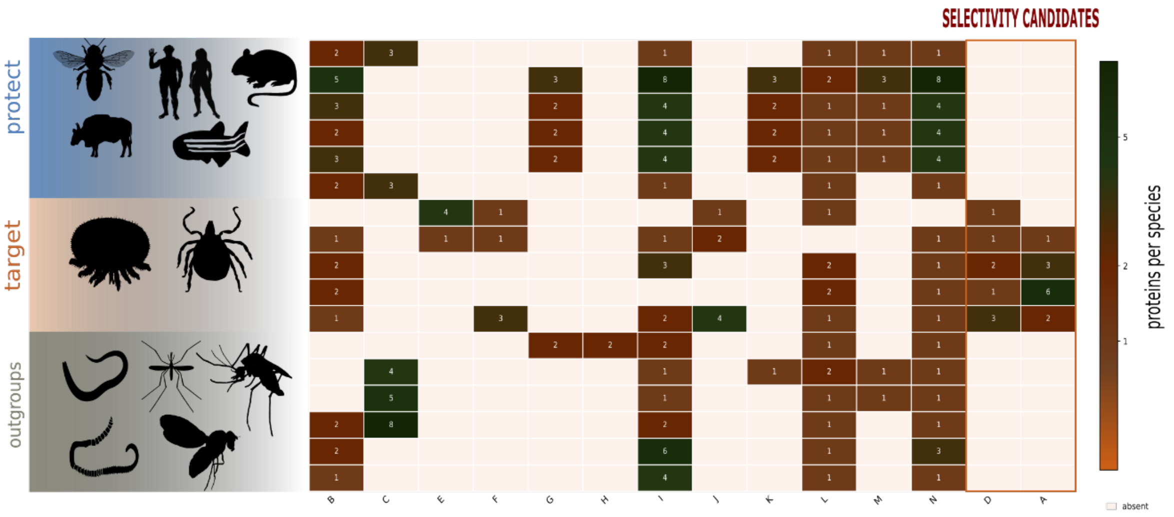


**Figure S2.** Selectivity profile of the 14 iGluR subfamilies across the 17 reference proteomes. Rows are species, grouped by role: protected (Apis mellifera, Bombus terrestris, Danio rerio, Homo sapiens, Bos taurus, Mus musculus), target (Rhipicephalus microplus, Ixodes scapularis, Tetranychus urticae, Sarcoptes scabiei, Varroa destructor) and outgroups (Caenorhabditis elegans, Aedes aegypti, Anopheles gambiae, Drosophila melanogaster, Lottia gigantea, Capitella teleta). Columns are subfamilies: numbers are proteins per species and the color is the same count on a log scale, with absence left blank. The two candidates, D and A, are moved to the right and boxed. Outgroups are drawn for context and take no part in the test. Subclade C, present only in the two bees among the protected species and holding eight Drosophila proteins, is the neuromuscular-junction group.

**Table S2.** The two selectivity candidates, the three further groups absent from every protected species, and the rejected Drosophila neuromuscular-junction group. Selectivity is the prevalence in the arachnids minus the prevalence in the protected species. q is the Benjamini–Hochberg corrected Fisher p-value over the 14 groups. The full table is in supplementary_case_4.tsv.

| Subclade | In arachnids | In protected | Selectivity | q | Copies per mite (proteins) |
|---|---|---|---|---|---|
| **D** | 5 / 5 | 0 / 6 | 1.00 | 0.030 | 1–3 (8) |
| **A** | 4 / 5 | 0 / 6 | 0.80 | 0.106 | 1–6 (12) |
| F | 3 / 5 | 0 / 6 | 0.60 | 0.212 | 1–3 (5) |
| J | 3 / 5 | 0 / 6 | 0.60 | 0.212 | 1–4 (7) |
| E | 2 / 5 | 0 / 6 | 0.40 | 0.509 | 1–4 (5) |
| C | 0 / 5 | 2 / 6 | −0.33 | 1.000 | 0 (0) |
